# Non-monotonic Relationship between Mutation Rate and Speed of adaptation under Antibiotic Exposure in *Escherichia coli*

**DOI:** 10.1101/2023.08.15.553341

**Authors:** Atsushi Shibai, Minako Izutsu, Hazuki Kotani, Chikara Furusawa

**Author notes:** Corresponding author (CF). These authors contributed equally to this work (Atsushi Shibai, Minako Izutsu).

## Abstract

The mutation is a fundamental source of biological evolution that create genetic variation in populations. Mutations can create new advantageous traits, but also potentially interfere with pre-existing organismal functions. Therefore, organisms may have evolved their mutation rates to appropriate levels to maintain or improve their fitness. In this study, we aimed to experimentally quantify the relationship between mutation rate and the speed of antibiotic resistance evolution. We conducted experimental evolution using twelve *Escherichia coli* mutator strains with increased mutation rates and five antibiotics. Our results showed that the highest mutation rate did not necessarily lead to the highest speed of adaptation, indicating a non-monotonic relationship between the speed of drug resistance evolution and mutation rate as expected. Moreover, this relationship was observed to differ among drugs, with significant differences in peak size observed between bacteriostatic and bactericidal antibiotics. We also successfully reproduced the mutation-rate dependence of the speed of adaptation using numerical simulation of a population dynamics model. These findings offer significant insights into the mutation rate’s evolution concomitant with the development of antibiotic resistance.

## Introduction

The mutation is a fundamental driving force behind biological evolution since it serves as a source of genetic variation within a population. Mutations have the ability to create new advantageous traits that are favored by natural selection while having a potential for interfering with pre-existing organismal functions. In other words, mutations exert a double-edged-sword influence on organisms, conferring both beneficial and deleterious effects [1], and that implies that mutation rates are adjusted to appropriate levels [1–3]. Specifically, under certain selection pressures, the speed of adaptive evolution can increase with the mutation rate, as demonstrated by several studies [4–6]. Accordingly, one would anticipate that alleles that change mutation rates can undergo positive selection in some conditions. For example, in the long-term experimental evolution of asexual *Escherichia coli* populations, cells with a significantly higher mutation rate called ‘mutator’ emerged [7,8], which likely acquired a larger number of beneficial mutations than the wild-type strain. However, when the mutation rate is too high, fitness is expected to decrease due to deleterious mutations that are more common than beneficial ones. Thus, under such circumstances, selection pressure can reduce the mutation rate. Indeed, several studies have demonstrated a reduction in mutation rate during evolution [3,9,10]. The beneficial and harmful effects of mutations imply a non-monotonic relationship between mutation rate and the speed of fitness increase.

Experimental evolution of asexual bacterial populations, aided by whole-genome sequencing, is a powerful tool for investigating the effects of mutation rates on evolution [1,3,4,6,11]. Bacterial strains with different mutation rates can be prepared, for example, by deleting genes related to DNA-repair mechanisms as the mutator strains. Using these strains, we can evaluate how changes in mutation rates affect the course of evolution. For example, Wagner and his colleagues conducted experimental evolution with engineered *E. coli* strains having four different mutation rates [12]. They evolved these strains for 3,000 generations in a minimal medium without explicit stressors. Although populations with higher mutation rates had greater genetic diversity, this diversity only benefited when the mutation rate was modestly high. The study demonstrated that the highest mutation rates they used were not optimal for evolution in the environment during the long-term cultivation, or stress tolerance in novel environments after evolution.

Although the question of how evolutionary dynamics depend on mutation rates is important and has been the focus of many studies, the extent to which this relationship is influenced by the selective environment remains uncertain. The relationship is expected to depend on multiple factors, including the frequency of beneficial and harmful mutations, population size, and selection pressure. Analyzing the mutation rate dependency of evolutionary dynamics experimentally allows us to capture the contributions of these factors, providing a better understanding of how mutation rates evolve in nature and laboratories. For example, the evolution of antibiotic resistance in microorganisms has been extensively studied in both laboratory and clinical settings [13–17]. Examining how evolutionary dynamics under antibiotics depend on mutation rates will provide valuable insight into the broader mechanisms underlying the emergence of antibiotic resistance.

In this study, we aimed to experimentally quantify the mutation rate dependency of the speed of antibiotic resistance evolution. To achieve this, we constructed twelve *E. coli* mutator strains with elevated mutation rates and conducted experimental evolution under five different antibiotics with varying action mechanisms. The results revealed a non-monotonic dependency between speed of adaptation, as quantified by the increasing rate of minimum inhibitory concentration (MIC), and the mutation rate. Furthermore, this dependency was found to differ between drugs, with significant differences between bacteriostatic and bactericidal antibiotics. We successfully reproduced these mutation-rate dependencies using numerical simulations of population dynamics model. This study provides valuable insights into the mechanisms that underlie the evolution of antibiotic resistance and highlights the importance of taking mutation rates into account when evaluating the efficacy of antibiotic treatments.

## Results

### Construction of hyper-mutable strains

We used the *E. coli* MDS42 strain as the wild-type (WT) and generated knockout mutants of the *mutS, mutH, mutL, mutT*, and *dnaQ* genes, which we denoted as S, H, L, T, and Q, respectively. The *mutS, mutH*, and *mutL* genes are involved in the mismatch repair machinery [18], *mutT* plays a role in maintaining replication fidelity [19], and *dnaQ* codes for the epsilon subunit of DNA polymerase III [20]. Deletion of these genes is known to cause a loss of replication fidelity and an increase in mutation rates. Additionally, we created seven double-gene-knockout strains and obtained 12 hyper-mutable strains, which are listed in Table 1.

**Table 1.** Summary of MA experiments. The two right-hand columns display synonymous (Syn) and non-synonymous (NSyn) base-pair substitutions (BPS), respectively, while the mean and standard deviation of three replicates are presented. A list of all detected mutations and an extended table showing the number of intergenic BPS and short Indel are shown in Supplementary tables S1 and S2, respectively.

| Strain | Abbr. | MA rounds | Generations | No. of Syn BPSs | No. of NSyn BPSs |
| --- | --- | --- | --- | --- | --- |
| MDS42 | WT | 66 | 1685.7 ± 7.0 | 0.67 ± 0.58 | 0.67 ± 0.58 |
| MDS42Δ*mutH* | H | 23 | 613.4 ± 1.7 | 4.33 ± 3.21 | 11.67 ± 1.53 |
| MDS42Δ*mutL* | L | 66 | 1675.5 ± 0.8 | 8.33 ± 2.31 | 15.00 ± 1.00 |
| MDS42Δ*mutS* | S | 66 | 1691.3 ± 9.3 | 11.00 ± 2.00 | 24.33 ± 1.15 |
| MDS42Δ*dnaQ* | Q | 66 | 1651.0 ± 5.9 | 8.67 ± 1.15 | 23.33 ± 2.89 |
| MDS42Δ*mutT* | T | 66 | 1679.7 ± 13.6 | 27.33 ± 2.31 | 157.67 ± 20.21 |
| MDS42Δ*mutL*Δ*mutH* | LH | 66 | 1651.4 ± 13.2 | 10.67 ± 1.53 | 15.00 ± 2.00 |
| MDS42Δ*mutS*Δ*mutH* | SH | 66 | 1691.1 ± 9.4 | 13.33 ± 5.13 | 21.00 ± 6.56 |
| MDS42Δ*mutL*Δ*dnaQ* | LQ | 62,62,70 | 1461.2 ± 96.0 | 545.67 ± 60.72 | 939.67 ± 81.5 |
| MDS42Δ*mutH*Δ*mutT* | HT | 23 | 610.8 ± 0.6 | 15.67 ± 5.03 | 76.33 ± 6.03 |
| MDS42Δ*mutL*Δ*mutT* | LT | 23 | 614.7 ± 2.5 | 16.00 ± 2.65 | 84.67 ± 14.57 |
| MDS42Δ*mutS*Δ*mutT* | ST | 23 | 605.0 ± 4.0 | 13.33 ± 4.51 | 54.33 ± 10.97 |
| MDS42Δ*dnaQ*Δ*mutT* | QT | 23 | 611.7 ± 2.6 | 13.33 ± 2.08 | 63.33 ± 7.09 |

To quantify the mutation rate, we conducted mutation accumulation (MA) experiments using the hyper-mutable strains as ancestors. Specifically, we propagated three lineages for each ancestor as single colonies on agar plate medium for 23-70 passages. We estimated the number of generations during the MA experiment by establishing a relationship between colony size and cell number [21–23]. Subsequently, we sequenced the samples at the end of the MA experiment to detect point mutations accumulated on the genome. The total number of identified base-pair substitutions (BPS) is summarized in Table 1.

We calculated the mutation rates per generation for each strain, following the previous study [24]. Initially, we determined the proportion of each pattern of synonymous base-pair substitution. Subsequently, we estimated the genome-wide mutation rate by dividing the number of accumulated mutations on a genome by the number of generations, and then normalizing it with the frequency of possible mutational patterns. As shown in Fig. 1, our results showed that the hyper-mutable strains exhibited mutation rates about 6 to 400 times higher than that of WT. The base-pair substitution patterns varied depending on the knockout gene(s), as shown in the pie charts in Fig. 1, which was consistent with previous studies [19,22,25–27]. Specifically, disruption of the mismatch-repair mechanism (*ΔmutS, ΔmutH, ΔmutL*) increased A:T to G:C and G:C to A:T substitutions, while knockout of the *mutT* gene increased only A:T to C:G. Additionally, for each hyper-mutable strain, we calculated dN/dS ratios, which represents the ratio of nonsynonymous to synonymous substitution rates. We found that dN/dS ratios did not differ from 1 for all strains except for WT, which had scarcely accumulated mutations (Fig. S1A). The dN/dS ratio close to 1 suggested that selection had little effect in our MA experiments.

**Figure 1.**
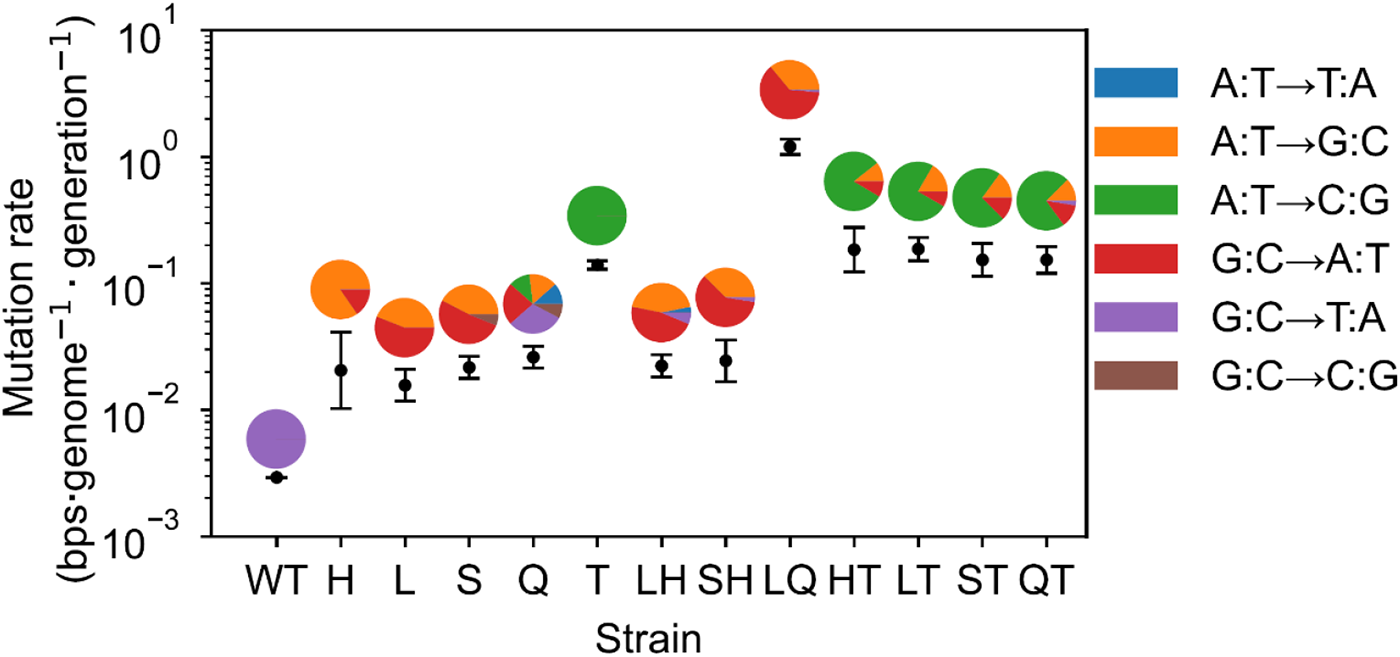
Mutation rates of hyper-mutable *E. coli* strains. Each dot with error bars represents the mean and standard deviation of the mutation rates observed in replicate MA lineages. The mutation rate was estimated using only synonymous mutations to exclude the effects of purifying selection. The pie charts above the dots show the distribution of substitution patterns identified in the hyper-mutable strains.

We then quantified the growth rates of the hyper-mutable strains in M9 minimum medium without adding any antibiotics. As shown in Fig. S1B, the growth rate decreased with increasing mutation rate, indicating deleterious effects of higher mutation rates.

### Quantifying speed of adaptation under antibiotics

To elucidate the relationship between mutation rate and evolutionary dynamics, we conducted experimental evolution of twelve hyper-mutable strains and the wild-type strain using five antibiotics with distinct action mechanisms, namely chloramphenicol (CP), trimethoprim (TP), amikacin (AMK), cefixime (CFIX), and ciprofloxacin (CPFX). We maintained four replica lines for each strain-antibiotic combination, resulting in a total of 260 individually evolving lines (13 strains × 5 antibiotics × 4 replicas). The cells were cultured in 200 µl of M9 medium supplemented with 20 amino acids (M9+AA medium) in a 96-well microtiter plate, to which each antibiotic was added as a two-fold dilution series (Fig. 2A). The cultivation began with a fixed initial cell concentration (OD_620_ value of 3×10^-4^, corresponding to approximately 2×10^5^ cells). After 24 hours of incubation, cells were collected from the well with the highest drug concentration among wells with OD values above a certain threshold (OD_620_=0.03). The collected cells were then transferred to a fresh medium containing the antibiotic dilution series, with the initial OD_620_ value of 3×10^-4^. We repeated this serial transfer procedure for nine days, corresponding to approximately 60 generations of cells.

**Figure 2.**
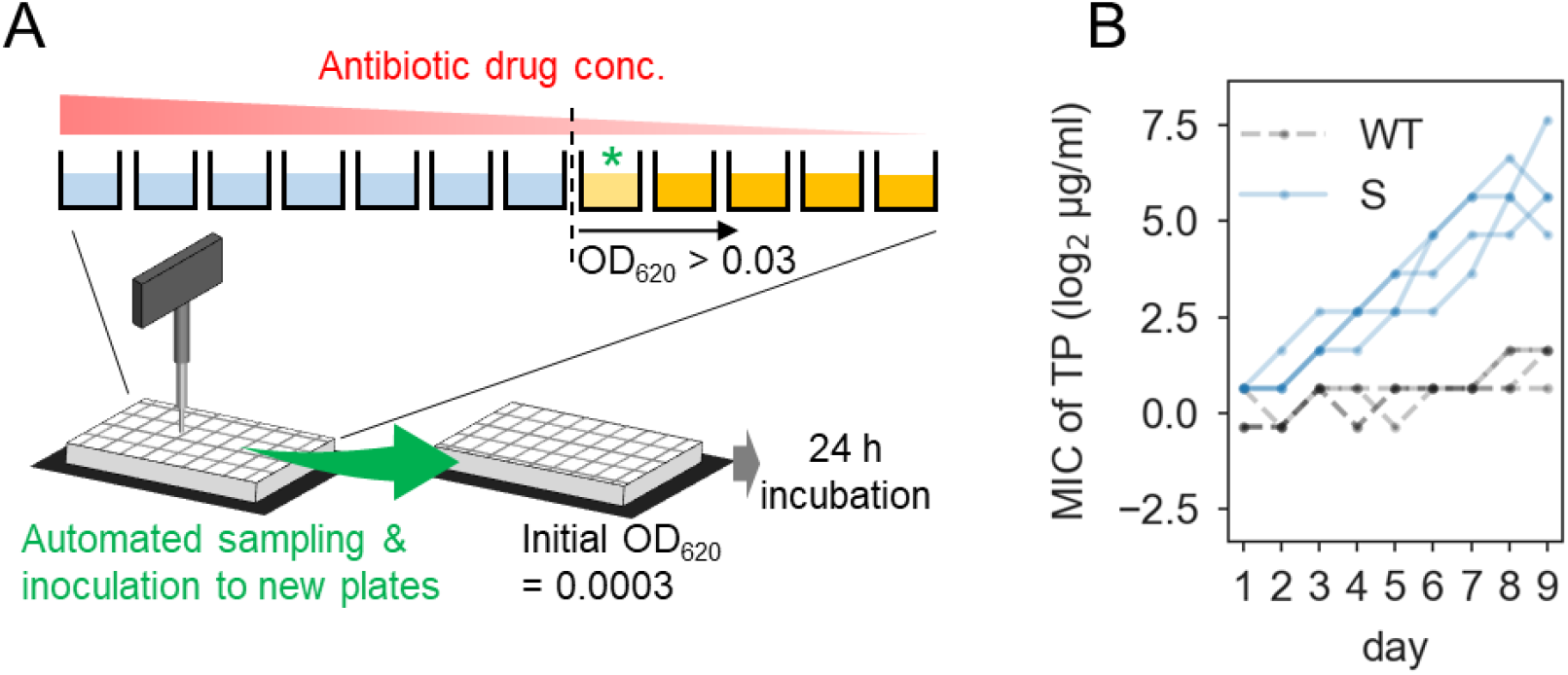
Experimental evolution under antibiotic selection pressure. (A) Schematic representation of the experimental procedure. A 96-well microtiter plate was prepared with two-fold serial dilutions of antibiotics. After 24 hours of cultivation, cells from the well with the highest drug concentration, in which the cell concentration (OD_620_) exceeded 0.03, were transferred to fresh medium with a drug concentration gradient. In this study, we designated the drug concentration of the selected well (green asterisk) as the minimal inhibitory concentration (MIC). The serial transfer cultures were iterated for 9 days. (B) Example of MIC changes during experimental evolution. The vertical axis shows the MIC for trimethoprim (TP). The black and blue lines represent the data for the WT and *ΔmutS* (S) strains, respectively. For each strain, data from four replicate series are overlaid. Data for all combinations of strains and drugs are presented in Fig. S2.

**Figure 3.**
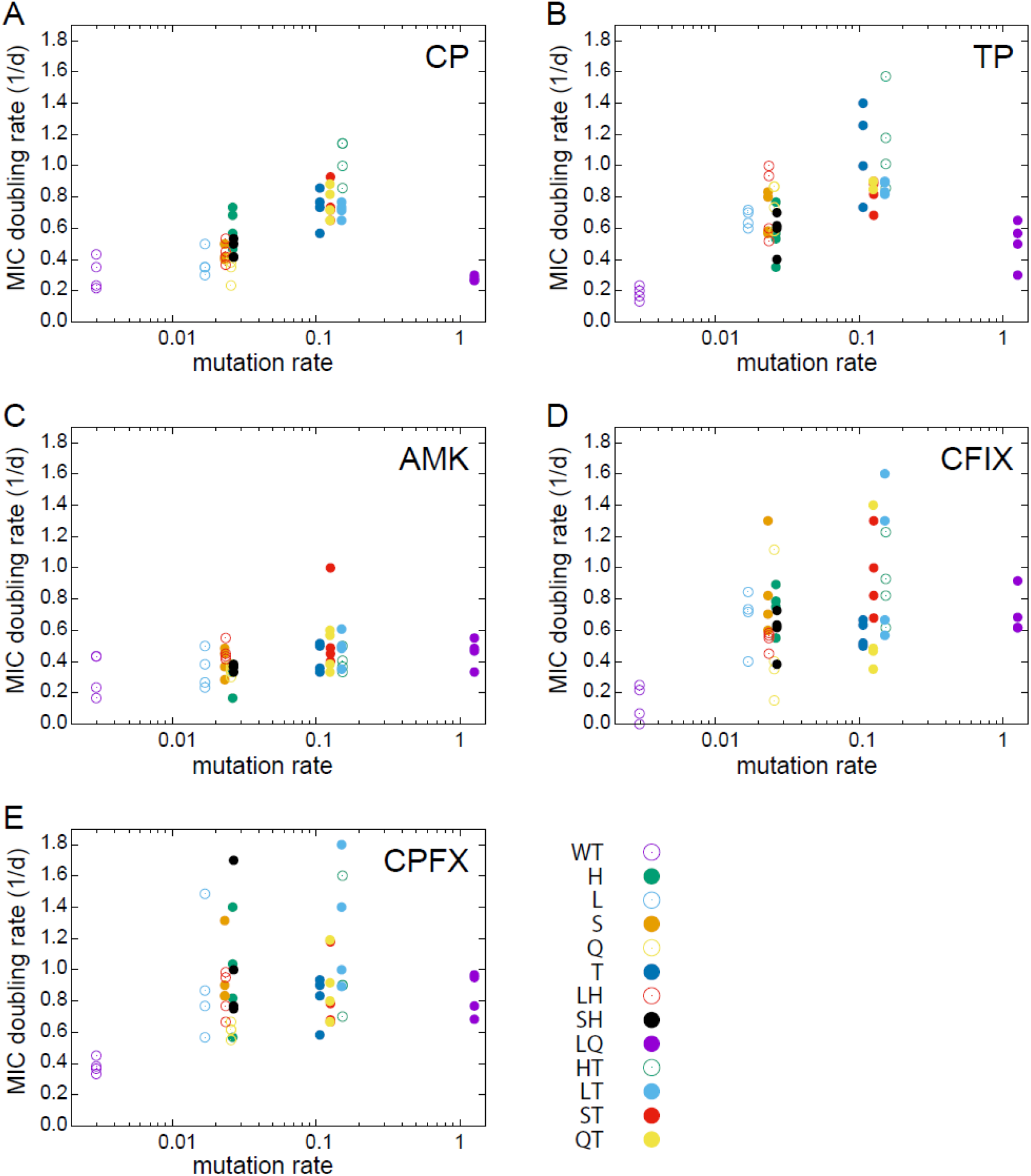
The relationship between mutation rate and MIC doubling rate. The dots represent experimental observations of 13 strains and four replicate serial transfer cultures.

In this study, we defined the minimum inhibitory concentration (MIC) as the drug concentration of the well from which cells were transferred. Fig. 2B shows typical examples of the time series of MIC under TP selection, starting from the wild-type (WT) and *ΔmutS* (S) strains, respectively (all MIC time series of 260 evolutionary lines are presented in Fig. S2). Using these time series data of MIC, we evaluated the speed of drug resistance evolution under antibiotic treatment by determining the doubling rate of the MIC per day, which was obtained using the linear fitting method. To assess the reproducibility of our observation, we conducted a replicate experiment over a shorter period of five days. The results indicated clear correlations between the speeds of adaptation in the replica series with different experimental periods (Fig. S3), demonstrating the reproducibility of the estimation of adaptation speed.

### Non-monotonical relationship between mutation rate and speed of adaptation

Figure 3 illustrates the relationship between mutation rate and speed of adaptation for each antibiotic. As can be seen, these dependencies do not always follow a monotonic pattern, wherein the strain LQ with the highest mutation rate often exhibited a smaller speed of adaptation than the strains with modest mutation rates. An interesting finding is that the mutation rate dependencies differ among the antibiotics used for selection. For CP and TP selections, there were significant decreases in MIC doubling rate at the highest mutation rate, while no significant decrease was observed for the other three drugs. CP and TP are known as bacteriostatic drugs, while AMK, CFIX, and CPFX are classified as bactericidal drugs. Thus, our results suggest that the effect of mutation rate on antibiotic resistance evolution depends on the action mechanisms of the drugs used for selection.

To elucidate the mutation rate dependency of adaptation speed, we employed a simple population dynamics model of multi-step resistance evolution that incorporates pharmacodynamic modeling [28,29]. In this model, *E. coli* populations can grow and increase their cell number up to a certain carrying capacity while accumulating mutations during their replication at a given mutation rate. Specifically, we considered the sequential accumulation of mutations that confer additive beneficial effects on growth under antibiotics (Fig. 4A). The population dynamics is described by the following deterministic differential equation:

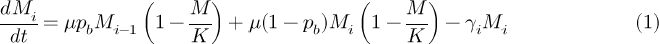

with 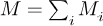. Here, *M_i_* represents the number of *E. coli* cells that has accumulated *i* beneficial mutations, *μ* is the growth rate, *γ_i_* is the death rate with *i* mutations, and *K* is the carrying capacity, respectively. The probability of acquiring a beneficial mutation, *p_b_*, is assumed to be proportional to the mutation rate, *p*, such that *p_b_*= *α_p_*, where *α* is the ratio of beneficial mutations to the total mutations. The first term on the right-hand side of Eq. (1) represents the influx of population caused by a single beneficial mutation. For simplicity, we do not allow reverse mutations that decrease the number of beneficial mutations.

By measuring the growth rate of hyper-mutable strains that we constructed, we observed that the growth rate *μ* decreases as the mutation rate *p* increases, as shown in Fig. S1, due to deleterious effects of high-mutation rate. We fit the data in Fig. S1 and obtained the following relationship: 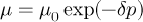 per hour, with *μ_0_* = 0.90 and *δ* = 0.54 . This relationship was used for the numerical simulations.

In our model, the addition of antibiotics results in an increase in the death rate of cells. However, the acquisition of beneficial mutations can enable cells to overcome the drug’s effects. We assume that the combined effect of antibiotics and beneficial mutations can be represented by the following death rate:

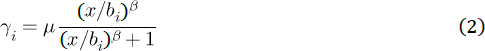

where *x* is log_2_-transformed concentration of antibiotics and the parameter *β* represents the sensitivity of the death rate to the addition of antibiotics (see Fig. 4B). At sufficiently high drug concentrations, the death rate *γ_i_* converges to the growth rate *μ*, and cells are unable to proliferate further. We do not consider the case where *μ<γ_i_*, as a subpopulation with *μ<γ_i_* is rapidly diluted in our serial transfer culture and becomes negligible. *b_i_*represents the benefit resulting from the accumulation of *i* mutations, which corresponds to the log_2_-transformed drug concentration that reduces the growth rate of the subpopulation by half (close to MIC in our experimental setting). In bacterial experimental evolution under antibiotics, MIC generally increases exponentially (see Fig. 2B for example). To model the exponential increase of MIC by mutations, we simply assume the following additive effect of mutations to the benefit: 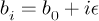, where *b_o_* and ϵ are constant parameters representing the resistance of non-mutated cell and benefit of single mutation, respectively. Note that the unit of benefit *b*_i_ is log-transformed concentration, thus the above additive effect of the mutations represents to an exponential increase in MIC. We simplify our model by neglecting the individuality of mutations and assuming that each mutation has the same beneficial effect and cost. Additionally, we neglect any positive or negative epistasis between mutations.

**Figure 4.**
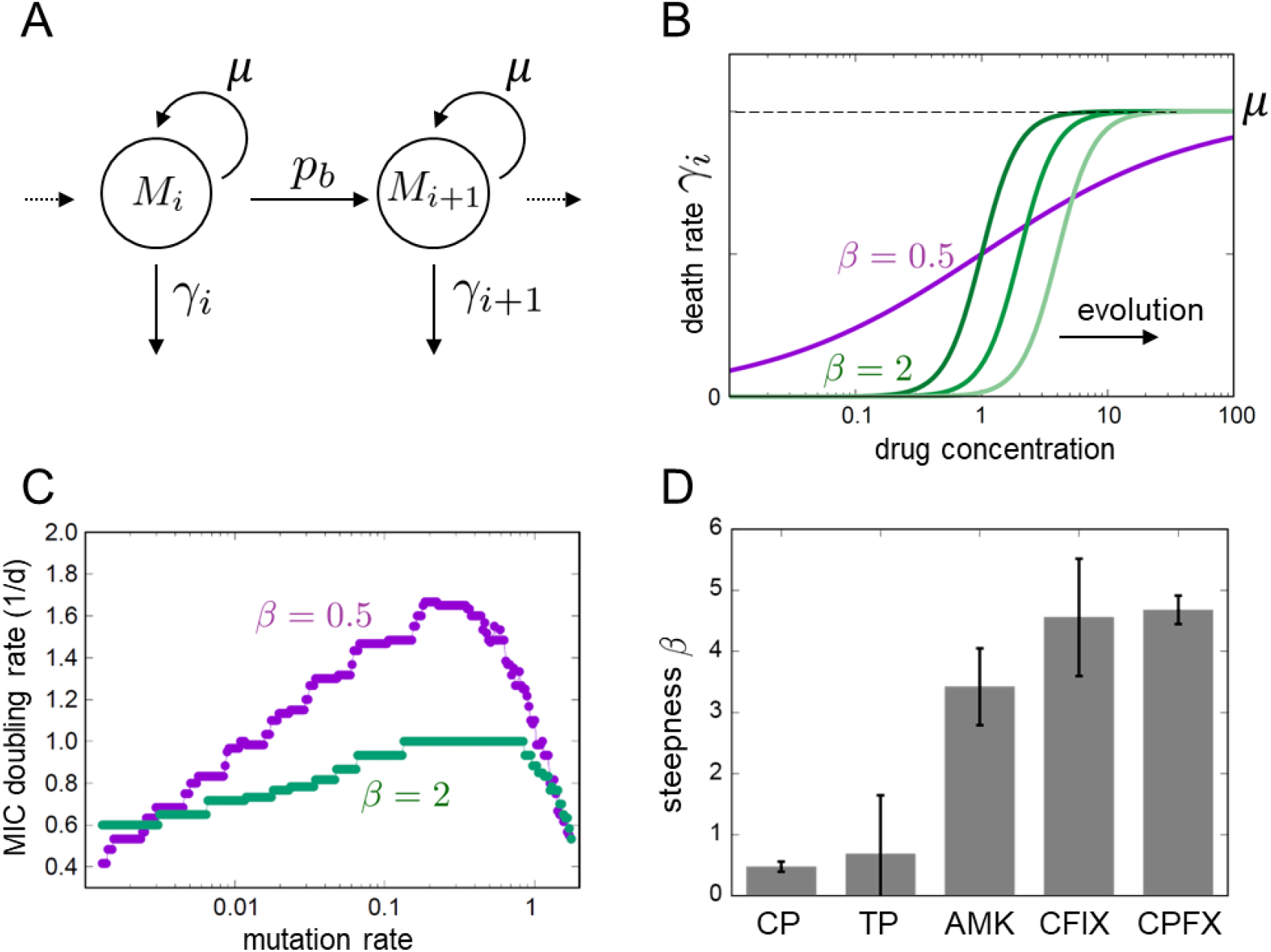
Parameter estimation by a population dynamics model. (A) Schematic representation of the model with multi-step resistance evolution. The mutated subpopulations (*M_i_*) with *i* mutations proliferate with the growth rate *μ*, mutate with the rate *p_b_*, and die at the death rate *γ_i_*. The death rate depends on the benefit of mutation *b_i_* conferred per mutation, as represented in Eq. (2). (B) The effect of antibiotics on the death rate. For the cases of *β* = 0.5 and *β* = 2, the death rates are plotted as a function of drug concentration, where *b_0_* is set to 1. The death rate curve shifts towards the right due to mutation accumulation, as depicted by the right green lines in the case of *β* = 2. (C) The simulated relationship between mutation rate and MIC doubling rate. The population dynamics model with mutation accumulation was utilized to simulate the increase in MIC observed in serial transfer experiments for the cases of *β* = 0.5 and *β* = 2. The parameters were set to *K* = 0.2, *b_0_* = 0.5, and ϵ = 0.5, respectively. (D) The estimated beta value for each drug. The mean value of β estimated through 100 bootstrap resampling is represented by the bars, while the error bars represent the standard error.

We utilized the population dynamics model to simulate the adaptive evolution of antibiotic resistance in our serial transfer setup (Fig. 2). In the initial condition, non-mutated cells were inoculated into a series of 2-fold dilutions of antibiotics with a constant initial density denoted as *M_0_* = 3 x 10^-4^ and *M_i_ = 0* for *i*≠ 0, in units of OD_620_. After 24 hours of cultivation, cells were sampled from an environment with highest drug concentration in which the total cell density exceeded a certain threshold (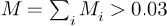), diluted, and subsequently transferred to a fresh environment for the next round of cultivation. Note that this procedure corresponds to our experimental design (Fig. 2A). This simulation procedure was iterated nine times, resulting in an increase in minimum inhibitory concentration (MIC) similar to that observed in Fig. 2B.

Figure 4C presents typical simulation results for the MIC doubling rate as a function of mutation rate. As shown in the figure, we observed that the curve is significantly influenced by *β*, which represents the sensitivity to antibiotics. When *β* is equal to 0.5, the MIC increases significantly with the increase in mutation rate before experiencing a drastic decrease when mutation rates become extremely high. Conversely, when the sensitivity is high, as indicated by the steeper kill curve in Fig. 4B, the change in the MIC with increasing mutation rates is smaller. This difference can be explained by the variation in the effective population sizes between these two conditions. In the case of the death rate with low steepness, cells with fewer beneficial mutations have a greater chance to grow, and these cells can also acquire further beneficial mutations. Therefore, with an increase in the mutation rate, the speed of adaptation can increase substantially. In contrast, when the sensitivity curve is steep, only a small fraction of cells with beneficial mutations can grow, and the MIC improvement is primarily governed by the time required for the mutated cells to take over the population. Hence, in this case, the effect of an increased mutation rate on the speed of adaptation is relatively small.

We used this population dynamics model to fit experimental data and elucidate the evolutionary dynamics presented in Fig. 5. The experimentally estimated values of *K*, *μ*_0_, and were used as fixed parameters, while ϵ, *b_0_*, and *β* were used as fitting parameters to minimize the residual sum of squares between experimental and simulated adaptation speeds for each antibiotic. Due to the difficulty in experimentally estimating the parameter *α*, representing the ratio of beneficial mutations, we assigned an arbitrary value and confirmed that the simulated results remained qualitatively unchanged within a certain range. The green solid lines in Fig. 5 represent the average of simulated results under the fitted parameters, with the light green range indicating the standard error estimated by bootstrap random resampling of experimental data.

**Figure 5.**
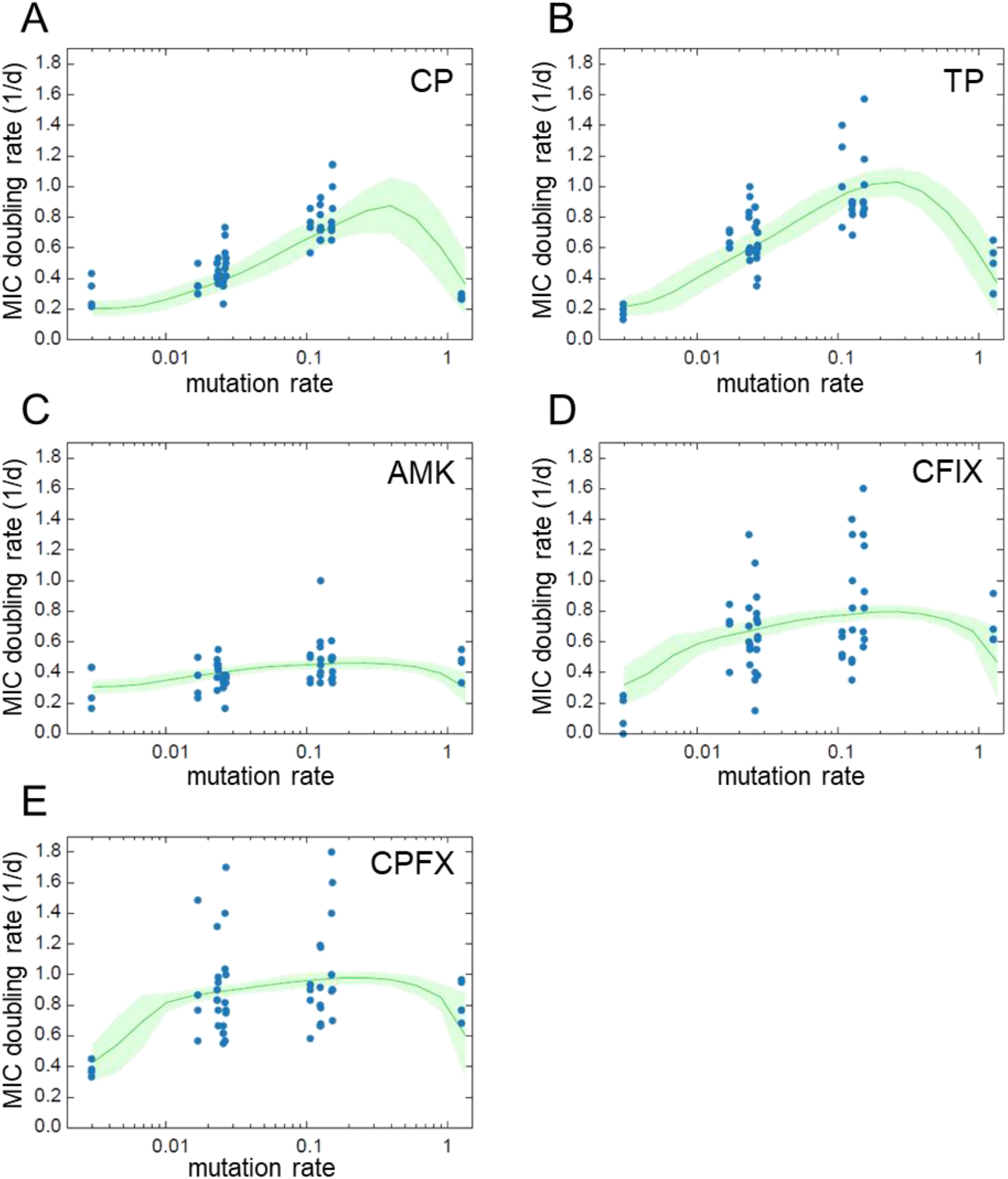
The fitting results of the population dynamics model to experimental observations. The blue dots represent experimental observation shown in Fig.3 and the green solid line shows the mean of simulated MIC changes generated with the parameters estimated from 100 bootstrap resampling, while the light green region represents the estimated standard error.

Our analysis revealed an interesting finding that the estimated sensitivity significantly differed among antibiotics (Fig. 4D), with relatively low values for CP and TP, and high values for the other three antibiotics. This result was consistent with the classification of bacteriostatic and bactericidal antibiotics, where the former is expected to have shallow killing curves. We were unable to find a clear interpretation for the estimated beneficial effect of each mutation ϵ (Fig. S4).

## Discussion

In this study, we conducted a quantitative analysis to investigate how changes in mutation rates affect the speed of antibiotic resistance evolution in *E. coli*. To achieve this, we generated a set of *E. coli* mutator strains by single and double deletions of genes related to error correction. We quantified the mutation rates of these mutators using MA experiments, resulting in a range of mutation rates caused by the gene deletions. We then performed experimental evolution starting from these mutators by serial transfer culture under five antibiotics, and we quantified the speed of adaptation by MIC increase rate over time. Our results revealed a non-monotonic curve of adaptation speed as a function of mutation rate (Fig. 3), with the peak sizes differing among the antibiotics used for selection. We explained the variation in the mutation rate dependency of the adaptation speed using numerical simulation using the simple population model.

Our findings indicated that the relationship between speed of adaptation and increasing mutation rates is dependent on the antibiotics used for selection. When using bacteriostatic drugs, a modest mutation rate led to a significant increase in speed of adaptation, while bactericidal drugs exhibited less dependency of adaptation speed on the mutation rate. This difference has provided novel insights into the conditions for the emergence of mutator strains. In cases where the slope of the killing curve for antibiotics is gradual, a greater number of cells have survived and obtain the opportunity to acquire beneficial mutations, resulting in stronger selection favoring higher mutation rates under drug treatment. Conversely, if the sensitivity to a drug shows a steep response, only a small fraction of cells survives and has a chance to acquire beneficial mutations. In this case, with a smaller effective population size, the benefit of increasing the mutation rate for resistance evolution is limited. While the relationship between mutation rate and the speed of adaptive evolution has been previously explored [5,12], these analyses were conducted under single selection environment. In contrast, the current study employed various drugs with distinct action mechanisms, implying that the effective population size, under the influence of strong drug selection, may impact the association between mutation rate and adaptation speed. Notably, our results indicate that the rate of adaptation under amikacin addition did not have a significant increase with increase in mutation rate. This result may shed light on the mechanism responsible for the infrequent emergence of amikacin-resistant strains [30]. As the emergence of mutator pathogens with multi-drug resistance is an increasingly important problem worldwide, our findings could serve as a basis for designing drug usage to suppress antibiotic resistance evolution.

As a recent advancement, Bollenbach and colleagues conducted drug-resistance evolution experiments with *E. coli* using an automated platform to quantify the evolvability of 98 strains with gene deletions [31]. Their study demonstrates epistasis between resistance mutations and genetic background by identifying gene deletion strains that affect the evolvability of drug resistance. The gene deletion library they used included *mutL* and *mutT*, which showed a modestly high mutation rate in our MA experiment. Additionally, they used Tetracycline and Chloramphenicol, which are both bacteriostatic drugs. Their experiment also showed that deletion of *mutL* and *mutT* generally accelerated the evolution of resistance to these drugs, which supports our conclusion. Note that they selected the gene deletions by ensuring that these deletions did not bring significant growth reduction to the cells in drug-free media. This gene selection contrasts with our research which focuses on the impact of higher mutation rates alongside growth reduction on evolvability. In this way, their study elaborates on the effect of gene deletions mainly involved in efflux pumps on drug resistance and resistance evolution, as well as the types of mutations that occur in the evolved strain. On the other hand, we shed light on the phenomenological behavior using quantitative parameters of mutation rate and growth rate, and the evolution of drug resistance. We found that qualitative changes in their phenomenological behavior appeared depending on whether the drug is bacteriostatic or bactericidal.

There are still questions regarding the relationship between mutation rate, fitness, and evolution of drug resistance. In our mutator strains, the highest mutation rate observed was about 1 base pair per genome per generation in the LQ strain. The LQ strain showed a significant reduction in growth rate, approximately 30% lower than the wild-type strain. Although the growth defect is likely due to the higher mutation rate, the mechanism for the growth rate reduction cannot always be attributed to the fixation of deleterious mutations. For example, assuming that 10% of all fixed mutations are lethal, the observed mutation rate in the LQ strain would only result in a 10% reduction in growth rate, despite the extremely high ratio of lethal mutations. Thus, fixation of deleterious or lethal mutations can only explain a small fraction of the growth defect, and unknown mechanisms remain for the growth defect under a high mutation rate. One possible mechanism is the cost associated with high-frequency gene repair, which could consume the cellular resources necessary for growth [24]. We anticipate that the elucidation of the growth defect will contribute to a better understanding of the evolution of mutation rates.

Evolution of mutation rates and effects of new mutations on reproductive success have been well studied [2,32] while pathogen/drug resistance has been used merely as marker traits to estimate mutation rates using the fluctuation test [33,34]. Given that recent studies have revealed drug resistance can be achieved by loss-of-function mutations [17] and studies that show the fitness cost of resistant genotypes [35,36], it is likely that the distribution of fitness effects in environments that contain drugs is qualitatively different from that studied under environments without drugs, and hypermutator genotypes might be favored by selection in the former environments. We provide the first empirical study that examines the relationship between the speed of drug-resistance evolution and mutation rates. Further experimental and theoretical investigations would help better understand the evolution of drug-resistant microbes as well as the emergence of drug-resistant cancer cells [37].

## Methods

### Bacterial strains and media

The insertion sequence (IS)-free *E. coli* K-12 substrain MDS42 [38], purchased from Scarab Genomics (Madison, Wisconsin, USA), was utilized as the wild-type (WT) strain in this study. Using this WT strain, we constructed 12 knockout strains of genes related to suppressing mutation rate (*mutS, mutH, mutL, mutT, dnaQ*) as listed in Table 1. These single- and double-gene knockouts were performed using the λ-Red homologous recombination method [39]. *E. coli* cells were cultured in modified M9 minimal medium supplemented with 20 amino acids (M9+AA medium) which includes 17.1 g/L Na_2_HPO_4_·12H_2_O, 3.0 g/L KH_2_PO_4_, 0.5 g/L NaCl, 2.0 g/L NH_4_Cl, 5.0 g/L glucose, 14.7 mg/L CaCl_2_·2H_2_O, 123.0 mg/L MgSO_4_·7H_2_O, 2.8 mg/L FeSO_4_·7H_2_O, and 10.0 mg/L thiamine hydrochloride (pH 7.0), and 20 canonical amino acids (0.02 mM Tyr and 0.05 mM of the remaining 19 amino acids) [40].

### Measurement of growth rate

To measure growth rate, cells precultured to yield an initial OD_600_ of 1×10^-4^ were inoculated into a 96-well microtiter plate containing 100 μL of M9+AA medium. The cells were then cultured at 34 °C with shaking at 300 rpm, and the cell density was measured at OD_600_ using an Infinite 200 PRO (TECAN) at 15-minute intervals. The specific growth rate was calculated based on three data points with minimum OD_600_ values under the condition that OD_600_ > 5×10^-3^.

### Mutation accumulation (MA) experiment

*E. coli* cells were subcultured as a single colony on M9+AA agar medium and incubated at 34 °C. To induce mutations, a colony was randomly selected and streaked onto a fresh agar medium every 2 days for a total of 23 or 62-70 passages. The number of generations per passage was estimated based on the colony diameter, which was previously shown to be a reliable indicator [21–23]. Specifically, we used a regression curve to estimate colony-forming units from the diameter and then calculated the number of generations based on the assumption that all the cells in a colony were derived from a single cell from the previous passage. The adopted relationship used in this study was (number of generations) = 24.51 + 2 log_2_ (colony diameter in mm) [23]. After each passage, we cultured the remaining cells of the selected colony in M9+AA liquid medium and stored them in glycerol. We repeated the experiment three times for each ancestral strain, resulting in a total of 39 MA lines.

### Whole-genome resequencing

Glycerol-stocked cells were grown in M9+AA medium until they reached full growth and were then harvested as a cell pellet using centrifugation. Genomic DNA was extracted from the cells using the DNeasy Blood & Tissue Kit (Qiagen). For library preparation prior to sequencing, the Nextera XT kit (Illumina) was used in a paired-end (2×300 bp) setting. The Illumina MiSeq platform was used to sequence the libraries with the MiSeq Reagent Kit v3, which provides 600 cycles (Illumina). The obtained short reads were mapped to the reference genome and point mutations accumulated during the MA experiment were detected using the breseq software [41].

### Experimental evolution under antibiotics

Cells were cultured in 200 µL volumes of M9+AA liquid medium in a 96-well microtiter plate. During the experiment, each culture line was exposed to 12 concentrations of antibiotics, corresponding to 11 wells with a two-fold dilution series and one drug-free well. The cell concentration was quantified by measuring the optical density at 620 nm (OD_620_). Cells were inoculated into these 12 wells at an initial OD_620_ value of 3×10^-4^. After 24 hours of cultivation at 34 °C with shaking at 300 rpm, cells were sampled from the well with the highest drug concentration among the wells that showed an OD_620_ value greater than 0.03. The sampled cells were diluted to an initial OD_620_ value of 3×10^-4^ and inoculated into fresh medium in 12 wells on a new plate, which was then incubated. For each combination of strain and antibiotics, four independent culture lines were maintained in parallel. The serial transfer experiment was conducted for nine days. The described experimental operations were performed using an automated culture system comprising Biomek NXP laboratory automation workstation (Beckman Coulter) in a clean booth, STX44 automated shaker incubator (LiCONiC), LPX220 plate hotel (LiCONiC), and FilterMax F3 microplate reader (Molecular Devices) [42].

### Parameter estimation

Custom C programs were used to implement the population dynamics model described in equation (1) and to optimize the model parameters. Specifically, a genetic algorithm was employed to estimate the values of the parameters, and *β*. The fitness function for the genetic algorithm was defined as the Euclidean distance between the experimentally observed MIC doubling rate and the rate obtained from simulations of the population dynamics model. The mean and standard errors of the parameters were estimated using bootstrap resampling of the experimental data.

## Data availability

The resequencing analysis data have been deposited at the DDBJ Sequence Read Archive (https://ddbj.nig.ac.jp/DRASearch/) under accession number DRA0163570.

## Acknowledgement

We thank Dr. Saburo Tsuru for fruitful discussions. This study was supported in part by JSPS KAKENHI (19K16114 and 21K15077 to A. S.; 19H05626 and 22K21344 to C.F.) and JST ERATO (JPMJER1902 to C.F.).

## Author contributions

Conceptualization: A.S., M.I. and C.F.; methodology: A.S., M.I., and C.F.; investigation: A.S., M.I., H.K., and C.F.; supervision: C.F; writing manuscript: A.S., M.I., and C.F.; funding acquisition: A.S. and C.F.; project administration: C.F.

---

**Figure S1.**
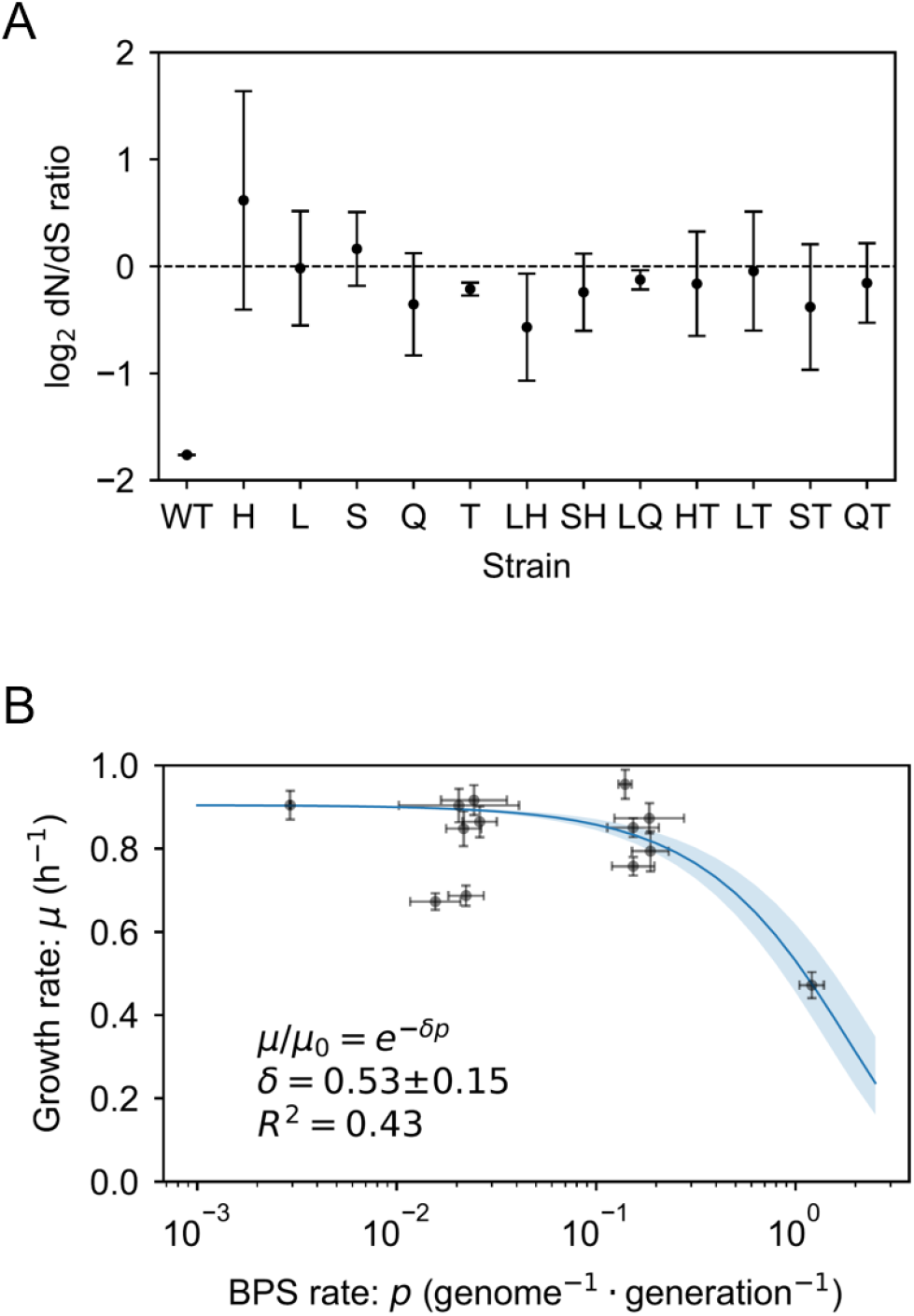
(A) Neutrality in mutation accumulation. The dN/dS ratio was calculated for each hyper-mutable strain and the wild-type strain. Error bars show the standard deviations between the MA lineages. (B) Relationship between growth rate and mutation rate. The horizontal error bars show the standard deviation across MA lineages, whereas the vertical error bars represent the standard deviation among replicate experiments in growth rate measurement. The sample sizes in the growth rate measurements were n=20 for wild-type strains and n=10 for mutant strains.

**Figure S2.**
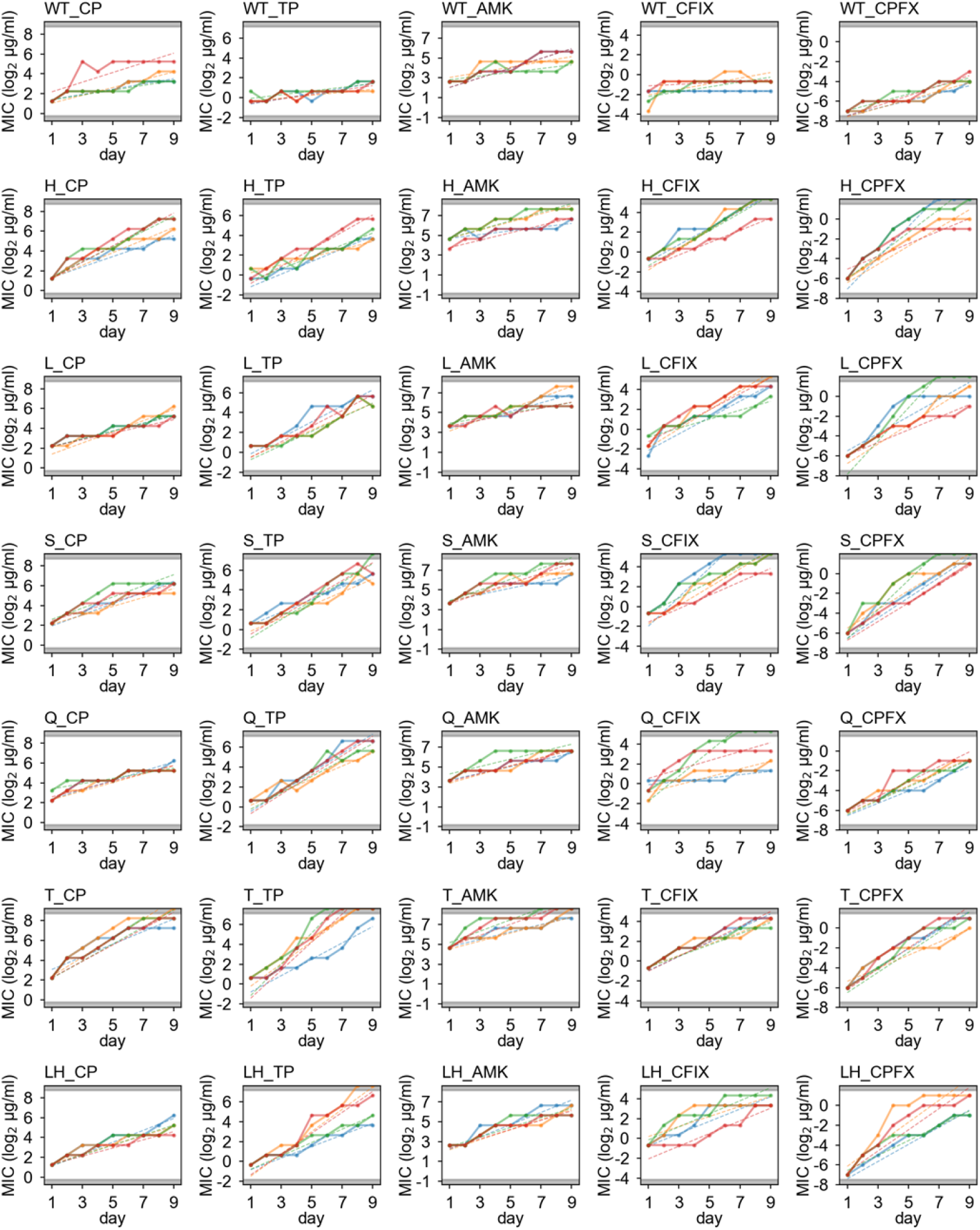
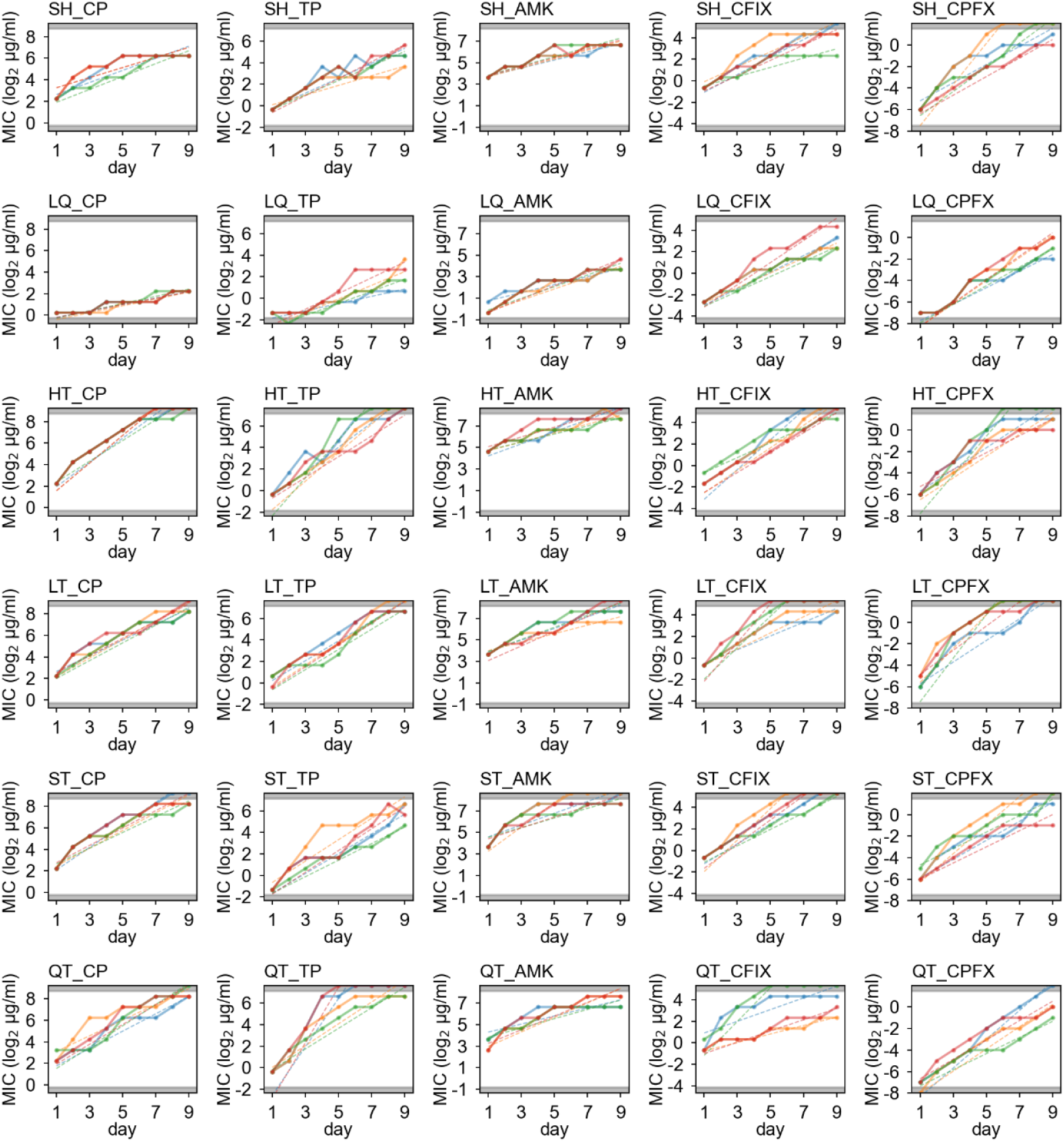
Experimental evolution of hyper-mutable strains under antibiotics. For all combinations of drugs and strains, the change of MIC over time are plotted. For each plot, data from four replicate series are overlaid. The dots and solid lines show experimental data, while dashed lines represent linear regressions of the increase in the evolutionary curve, performed via the least-squares method. To focus the regression analysis solely on the ascendant portion of the curve, data points indicating the minimum and maximum MIC values achievable within our experimental parameters (highlighted by gray fills) were systematically excluded.

**Figure S3.**
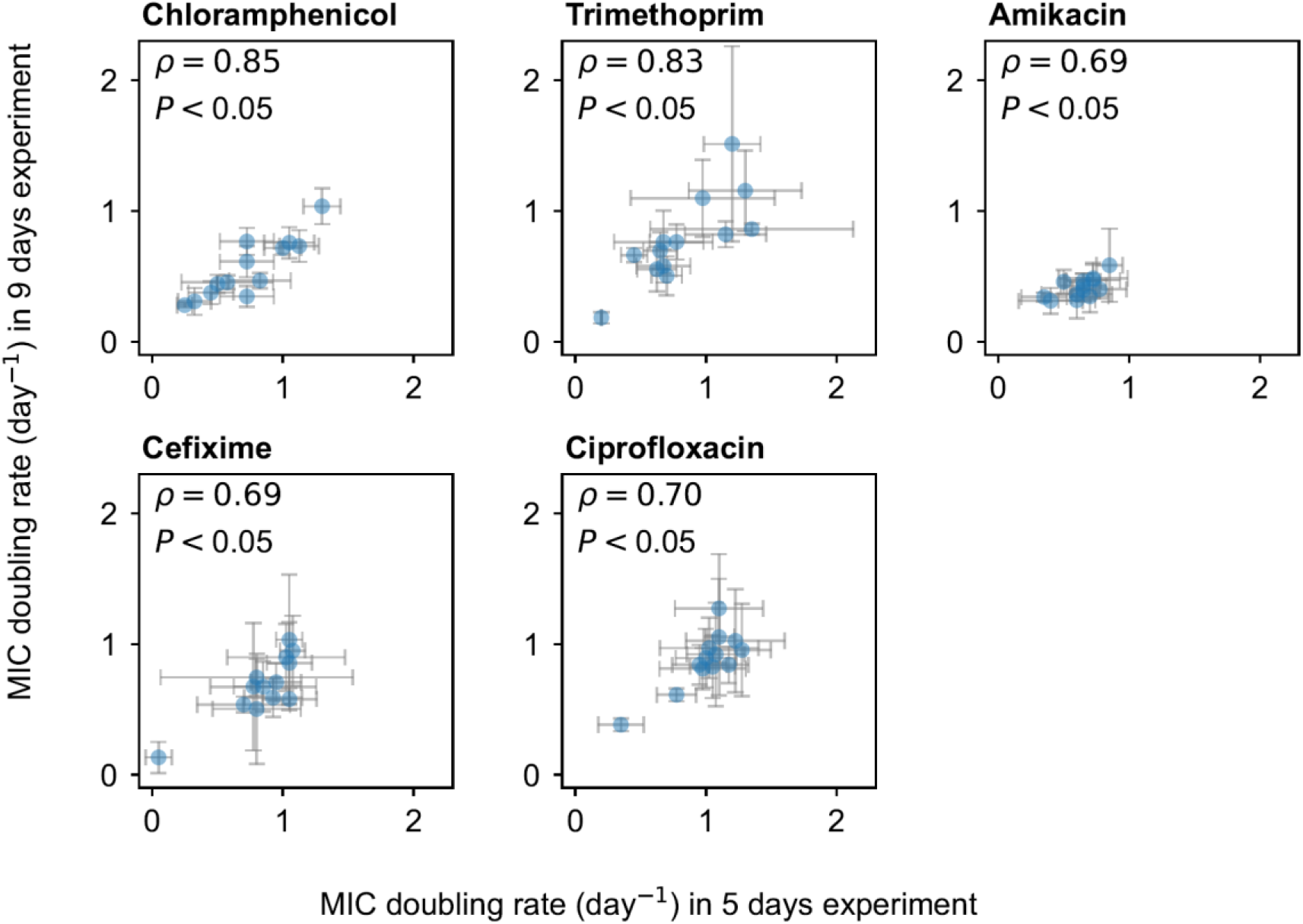
Reproducibility of the adaptation speed quantification. The MIC doubling rates were estimated by conducting independent experimental evolution trials with varying duration (9 days and 5 days, respectively). Each dot and error bars show the mean and standard deviation of the slope of the evolutionary curve, respectively, fitted per replicate for each strain as illustrated in Fig. S2. The Spearman’s rank correlation coefficient and corresponding P-value were computed for these data points (N=13).

**Figure S4.**
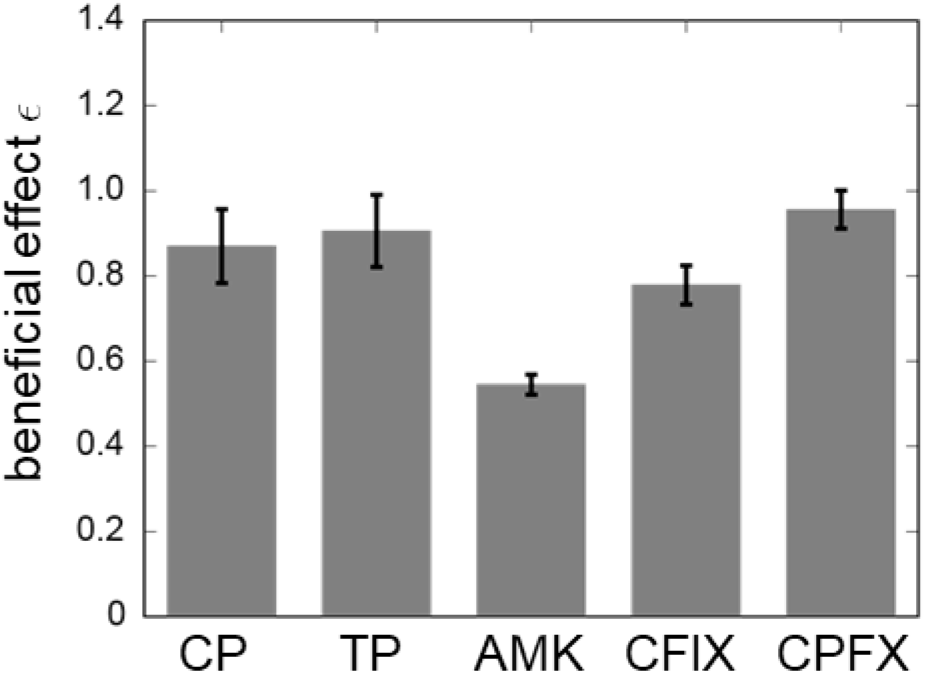
Estimated parameter representing the beneficial effect of each mutation. The mean value estimated through 100 bootstrap resampling is represented by the bars, while the error bars represent the standard error.

